# The COVID-19 PHARMACOME: A method for the rational selection of drug repurposing candidates from multimodal knowledge harmonization

**DOI:** 10.1101/2020.09.23.308239

**Authors:** Bruce Schultz, Andrea Zaliani, Christian Ebeling, Jeanette Reinshagen, Denisa Bojkova, Vanessa Lage-Rupprecht, Reagon Karki, Sören Lukassen, Yojana Gadiya, Neal G. Ravindra, Sayoni Das, Shounak Baksi, Daniel Domingo-Fernández, Manuel Lentzen, Mark Strivens, Tamara Raschka, Jindrich Cinatl, Lauren Nicole DeLong, Phil Gribbon, Gerd Geisslinger, Sandra Ciesek, David van Dijk, Steve Gardner, Alpha Tom Kodamullil, Holger Fröhlich, Manuel Peitsch, Marc Jacobs, Julia Hoeng, Roland Eils, Carsten Claussen, Martin Hofmann-Apitius

## Abstract

The SARS-CoV-2 pandemic has challenged researchers at a global scale. The scientific community’s massive response has resulted in a flood of experiments, analyses, hypotheses, and publications, especially in the field of drug repurposing. However, many of the proposed therapeutic compounds obtained from SARS-CoV-2 specific assays are not in agreement and thus demonstrate the need for a singular source of COVID-19 related information from which a rational selection of drug repurposing candidates can be made. In this paper, we present the COVID-19 PHARMACOME, a comprehensive drug-target-mechanism graph generated from a compilation of 10 separate disease maps and sources of experimental data focused on SARS-CoV-2 / COVID-19 pathophysiology. By applying our systematic approach, we were able to predict the synergistic effect of specific drug pairs, such as Remdesivir and Thioguanosine or Nelfinavir and Raloxifene, on SARS-CoV-2 infection. Experimental validation of our results demonstrate that our graph can be used to not only explore the involved mechanistic pathways, but also to identify novel combinations of drug repurposing candidates.

## Introduction and Motivation

COVID-19 is the term coined for the pandemic caused by SARS-CoV-2. Unprecedented in the history of science, this pandemic has elicited a worldwide, collaborative response from the scientific community. In addition to the strong focus on the epidemiology of the virus^1 2 3^, experiments aimed at understanding mechanisms underlying the pathophysiology of the virus have led to new insights in a comparably short amount of time^4 5 6 7^.

In the field of computational biology, several initiatives have started generating disease maps that represent the current knowledge pertaining to COVID-19 mechanisms^8 9 10 11^. Such disease maps have proven valuable before in diverse areas of research such as ^12 13 14 15^.

When taken together with related work including cause-and-effect modeling^8^, entity relationship graphs^16^, and pathways^17^; these disease maps represent a considerable amount of highly curated “knowledge graphs” which focus primarily on COVID-19 biology. Here, we use the term “mechanism” to describe a single, or multiple cause-and-effect relationships (i.e. a subgraph), “pathways” to refer to a well-established series of interactions resulting in cellular change or a defined product, and “models” for describing a collection of experimental data or known interactions defined in the context of a particular biological process or pathology. As of July 2020, a collection consisting of 10 models representing core knowledge about the pathophysiology of SARS-CoV-2 and its primary target, the lung epithelium, was shared with the public.

With the rapidly increasing generation of data (e.g. transcriptome^18^, interactome^19^, and proteome^20^ data), we are now in the position to challenge and validate these COVID-19 pathophysiology knowledge graphs with experimental data. This is of particular interest as validation of these knowledge graphs bears the potential to identify those disease mechanisms highly relevant for targeting in drug repurposing approaches.

The concept of drug repurposing (the secondary use of already developed drugs for therapeutic uses other than those they were designed for) is not new. The major advantage of drug repurposing over conventional drug development is the massive decrease in time required for development as important steps in the drug discovery workflow have already been successfully passed for these compounds^21 22^.

Our group and many others have already begun performing assays to screen for experimental compounds and approved drugs to serve as new therapeutics for COVID-19. Dedicated drug repurposing collections, such as the Broad Institute library^23^, and the even more comprehensive ReFRAME library^24^, were used to experimentally screen for either viral proteins as targets for functional inhibition^25^, or for virally infected cells in phenotypic assays^26^. In our own work, compounds were assessed for their inhibition of virus-induced cytotoxicity using the human cell line Caco-2 and a SARS-CoV-2 isolate^27^. A total of 63 compounds with IC50 < 20 μM were identified, from which 90% have not yet been previously reported as being active against SARS-CoV-2. Out of the active compounds, 31 are approved drugs, 23 are in phases 1-3 and 9 are preclinical candidate molecules. The described mechanisms of action for the inhibitors included kinase signaling, PDE activity modulation, and long chain acyl transferase inhibition (e.g. “azole class antifungals”).

The approach presented here integrates experimental results and the output from other informatic pipelines, and combines proprietary and public data to provide a comprehensive overview on the therapeutic efficacy of candidate compounds, the mechanisms targeted by these candidate compounds, and a rational approach to test the drug-mechanism associations for their potential in combination therapy.

## Methodology

### Generation of the COVID-19 PHARMACOME

Disparate COVID-19 disease maps focus on different aspects of COVID-19 pathophysiology. Based on comparisons of the COVID-19 knowledge graphs, we found that not a single disease map covers all aspects relevant for the understanding of the virus, host interaction and the resulting pathophysiology. Thus, we optimized the representation of essential COVID-19 pathophysiology mechanisms by integrating several public and proprietary COVID-19 knowledge graphs, disease maps, and experimental data (Supplementary Table 1) into one unified knowledge graph, the COVID-19 *Supergraph*.

To this end, we converted all knowledge graphs and interactomes into OpenBEL^28^, a language that is both ideally suited to capture and to represent “cause-and-effect” relationships in biomedicine and is fully interoperable with major pathway databases^29 30^. In order to ensure that molecular interactions were correctly normalized, individual pipelines were constructed for each model to convert the raw data to the OpenBEL format. For example, the COVID-19 Disease Map contained 16 separate files, each of which represented a specific biological focus of the virus. Each file was parsed individually and the entities and relationships that did not adhere to the OpenBEL grammar were mapped accordingly. Whilst most of the entities and relationships in the source disease maps could be readily translated into OpenBEL, a small number of triples from different source disease maps required a more in-depth transformation. When classic methods of naming objects in triples failed, the recently generated COVID-19 ontology^31^ as well as other available standard ontologies and vocabularies were used to normalize and reference these entities.

In addition to combining the listed models, we also performed a dedicated curation of the COVID-19 supergraph in order to annotate the mechanisms pertaining to selected targets and the biology around prioritized repurposing candidates. The resulting BEL graphs were quality controlled and subsequently loaded into a dedicated graph database system underlying the Biomedical Knowledge Miner (BiKMi), which allows for comparison and extension of biomedical knowledge graphs (see http://bikmi.covid19-knowledgespace.de).

Once the models were converted to OpenBEL and imported into the database, the resulting nodes from each mechanism-based model were compared (Figure 1). Even when separated by data origin type, the COVID-19 knowledge graphs had very little overlap (3 shared nodes between all manually curated models and no shared nodes between all models derived from interaction databases), but by unifying the models, our COVID-19 supergraph improves the coverage of essential virus- and host-physiology mechanisms substantially.

**Figure 1:**
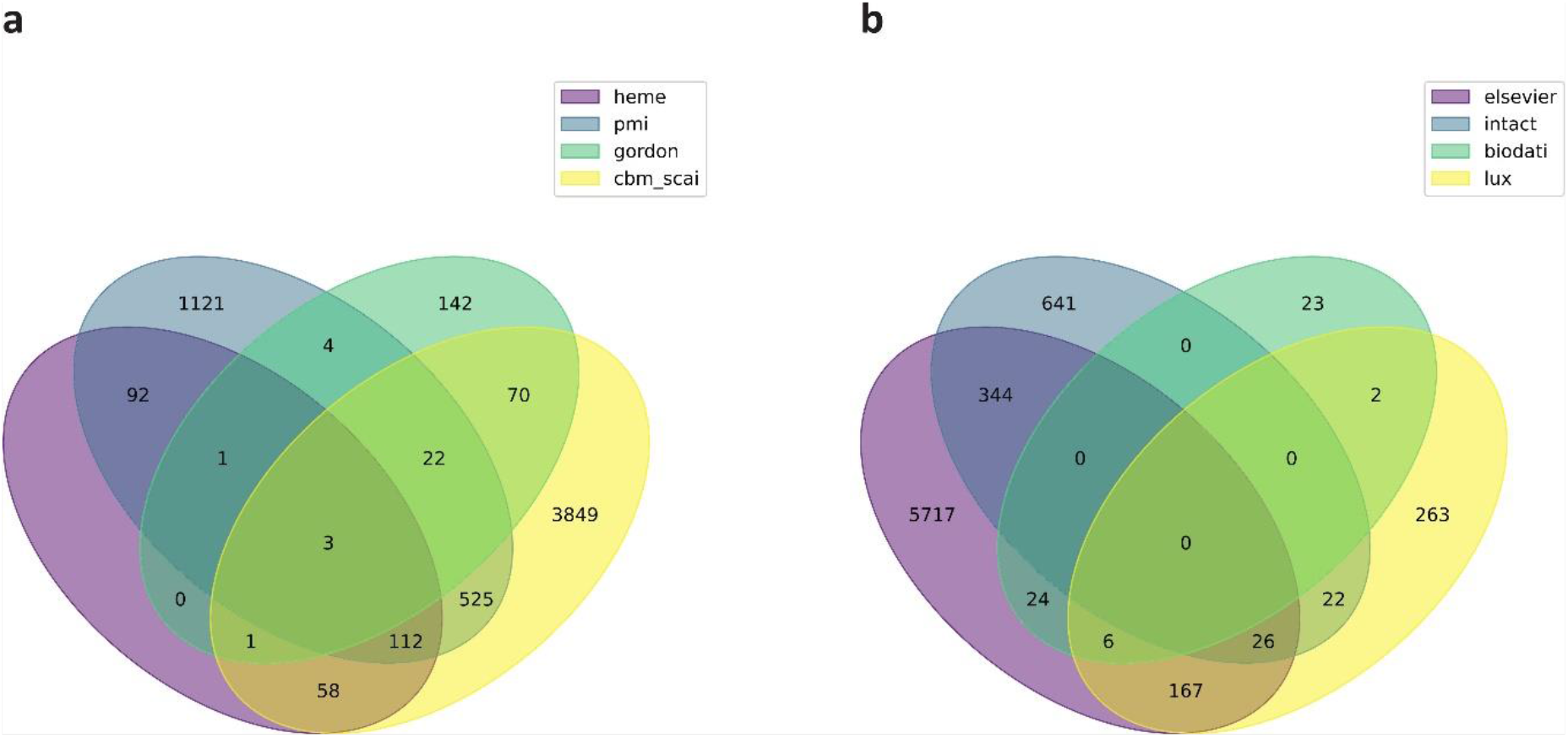
Venn diagrams comparing major mechanistic models in the COVID-19 supergraph. Mechanism-based models were divided, and their entities compared within their resulting subgroups. Model abbreviations are defined in Supplementary Table 1. a) Manual node comparison shows the overlap of entities in the models that are knowledge-based, manually curated relationships that have been directly encoded in OpenBEL. b) Automated node comparison shows the overlap of entities in models re-encoded into OpenBEL from other formats (e.g. SBML models).

Additionally, by enriching the COVID-19 supergraph with drug-target information linked from highly curated drug-target databases (DrugBank, ChEMBL, PubChem), we created an initial version of the **COVID-19 PHARMACOME**, a comprehensive drug-target-mechanism graph representing COVID-19 pathophysiology mechanisms that includes both drug targets and their ligands (Figure 2). In order to maximize its utility, this network includes both experimentally validated drug-target relationships as well as a wide distribution of biological entities and concepts (Supplementary Figure 1). The entire COVID-19 PHARMACOME was manually inspected and re-curated; this graph database is openly accessible to the scientific community at http://graphstore.scai.fraunhofer.de.

**Figure 2:**
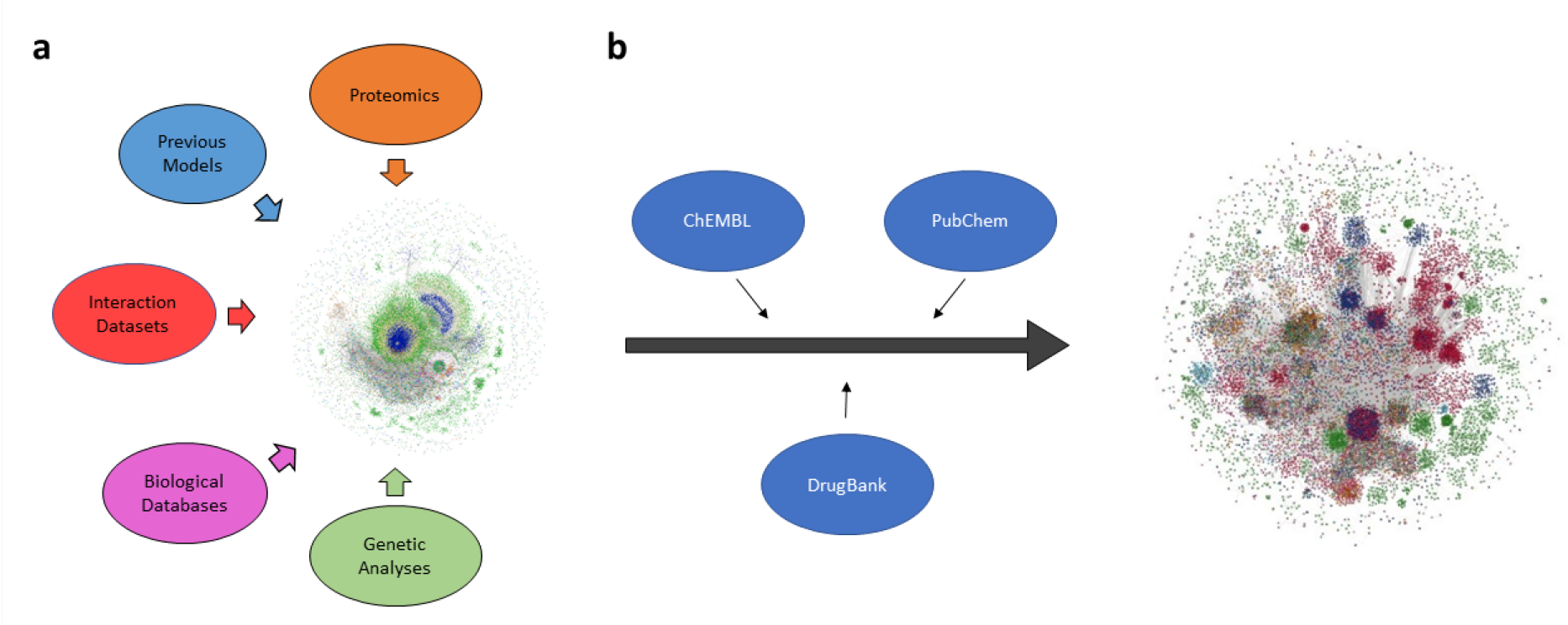
The COVID-19 supergraph integrates drug-target information to form the COVID-19 PHARMACOME. a) An aggregate of 10 constituent COVID-19 computable models covering a wide spectrum of pathophysiological mechanisms associated with SARS-CoV-2 infection or harmonized to generate the mechanism-based COVID-19 supergraph. b) The COVID-19 supergraph is annotated with drug-target information from a variety of curated sources to generate the COVID-19 PHARMACOME composed of 150662 nodes (representing proteins, pathologies, and other biological entities/concepts) and 573929 edges (indicating relationships or interactions between the pair of nodes they connect).

### Systematic review and integration of information from phenotypic screening

At the time of the writing of this paper, six phenotypic cellular screening experiments have been shared via archive servers and journal publications (Supplementary Table 2). Although only a limited number of these manuscripts have been officially accepted and published, we were able to extract their primary findings from the pre-publication archive servers. A significant number of reports on drug repurposing screenings in the COVID-19 context demonstrate how appealing the concept of drug repurposing is as a quick answer to the challenge of a global pandemic. Drug repurposing screenings were all performed with compounds for which a significant amount of information on safety in humans and primary mechanism of action is available. We generated a list of “hits” from cellular screening experiments while results derived from publications that reported on *in-silico* screening were ignored. Therefore, we keep a strict focus on well-characterized, well-understood candidate molecules in order to ensure that one of the pivotal advantages of this knowledge base is its use for drug repurposing.

### Subgraph annotation

The COVID-19 PHARMACOME contains several subgraphs, three of which correspond to major views on the biology of SARS-CoV-2 as well as the clinical impact of COVID-19:

- the ***viral life cycle*** subgraph focuses on the stages of viral infection, replication, and spreading.
- the ***host response*** subgraph represents essential mechanisms active in host cells infected by the virus.
- the ***clinical pathophysiology*** subgraph illustrates major pathophysiological processes of clinical relevance.

These subgraphs were annotated by identifying nodes within the COVID-19 PHARMACOME that represent specific biological processes or pathologies associated with each subgraph category and traversing out to their first-degree neighbors. For example, a biological process node representing “viral translation” would be classified as a starting node for the viral life cycle subgraph while a node defined as “defense response to virus” would be categorized as belonging to the host response subgraph. Though the viral life cycle and host response subgraphs contain a wide variety of node types, the pathophysiology subgraph is restricted to pathology nodes associated with either the SARS-CoV-2 virus or the COVID-19 pathology.

### Mapping of gene expression data onto the COVID-19 PHARMACOME

Two single cell sequencing data sets representing infected and non-infected cells directly derived from human samples^32^ and cultured human bronchial epithelial cells^33^ (HBECs) were used to identify the areas of the COVID-19 PHARMACOME responding at gene expression level to SARS-CoV-2 infection. Details of the gene expression data processing and mapping are available in the supplementary material (section gene expression data analysis).

### Pathway enrichment

Associated pathways for subgraphs and significant targets were identified using the Enrichr^34^ feature of the gseapy Python package^35^. Briefly, gene symbol lists were assembled from their respective subgraph or dataset and compared against multiple pathway gene set libraries including Reactome, KEGG, and WikiPathways. To account for multiple comparisons, *p*-values were corrected using the Benjamini-Hochberg^36^ method and results with *p*-values < 0.01 were considered significantly enriched.

### Drug repurposing screening

We performed phenotypic assays to screen for repurposing drugs that inhibit the replication and the cytopathic effects of virus infection. A derivative of the Broad repurposing library was used to incubate Caco-2 cells before infecting them with an isolate of SARS-CoV-2 (FFM-1 isolate, see ^37^). Survival of cells was assessed using a cell viability assay and measured by high-content imaging using the Operetta CLS platform (PerkinElmer). Details of the drug repurposing screening are described in the supplemental material.

### Drug combinations assessment with anti-cytopathic effect measured in Caco-2 cells

As described in Ellinger et al.,^38^ we challenged four combinations of five different compounds with the SARS-CoV-2 virus in four 96-well plates containing two drugs each. Eight drug concentrations were chosen ranging from 20 μM to 0.01 μM, diluted by a factor of 3 and positioned orthogonally to each other in rows and columns. No pharmacological control was used, only cells with and without exposure to SARS CoV-2 virus at 0.01 MOI.

In addition, recently published data from the work of Bobrowski et al.^39^, were mapped to the COVID-19 PHARMACOME and compared to the results of the combinatorial treatment experiments performed here.

## Results

### Comparative analysis of the hits from different repurposing screenings

Data from six published drug repurposing screenings were downloaded, and extensive mapping and curation was performed in order to harmonize chemical identifiers. The curated list of drug repurposing “hits” together with an annotation of the assay conditions is available under http://chembl.blogspot.com/2020/05/chembl27-sars-cov-2-release.html

Initially, we analyzed the overlap between compounds identified in the reported drug repurposing screening experiments. Figure 3A shows no overlap between experiments, which is not surprising, as we are comparing highly specific candidate drug experiments with screenings based on large drug repositioning libraries. However, the overlap is still quite marginal for those screenings where large compound collections (Broad library, ReFRAME library) have been used.

**Figure 3:**
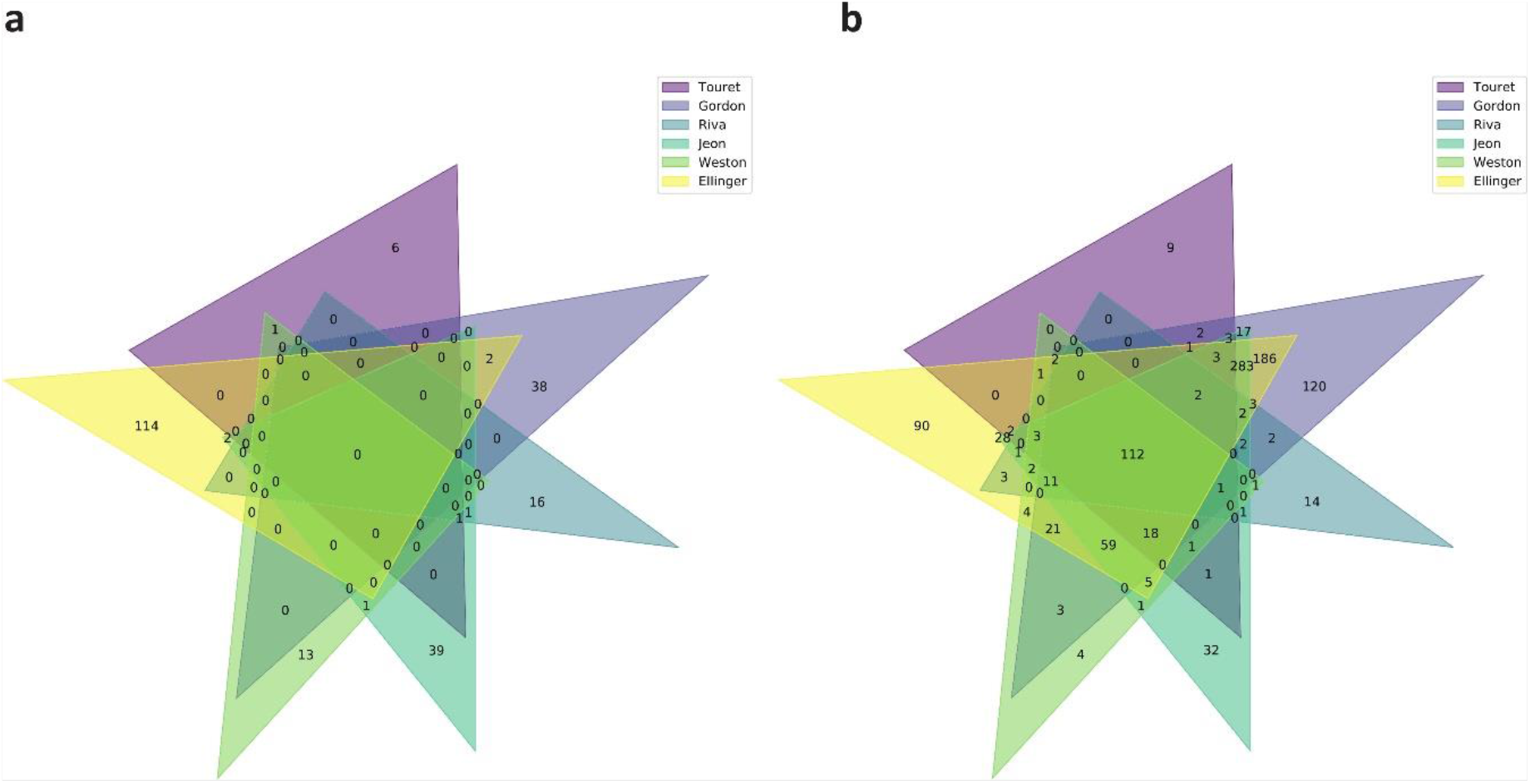
Overlap of compound hits between different drug repurposing screening experiments. a) Direct comparison of overlapping hits in drug repurposing screenings revealed no overlap between the experiments. These experiments were performed using different cell types (Vero E6 cells and Caco2 cells). b) Protein target space overlap between different COVID-19 drug repurposing screenings. Drug targets were identified by confidence level >= 8 and single protein targets according to the ChEMBL database. Comparison of experiments indicates over one hundred common protein targets.

### Mapping of repurposing hits to target proteins

In order to identify which proteins are targeted by the repurposing hits, and to investigate the extent to which there are overlaps between repurposing experiments at the target/protein level, we mapped all the identified compounds from the drug repurposing experiments to their respective targets. As most drugs bind to more than one target, we increase the likelihood of overlaps between the drug repurposing experiments when we compare them at the protein/target space. Indeed, Figure 3B shows an overlap of 112 targets between all the drug repurposing experiments, thereby creating a list of potential proteins for therapeutic intervention when the compound targets are considered rather than the compounds themselves.

### The COVID-19 PHARMACOME associates pathways derived from drug repurposing targets with pathophysiology mechanisms

A non-redundant list of drug repurposing candidate molecules that display activity in phenotypic (cellular) assays was generated and mapped to the COVID-19 PHARMACOME. Figure 4 shows the distribution of repurposing drugs in the COVID-19 cause-and-effect graph, the “responsive part” of the graph that is characterized by changes in gene expression associated with SARS-CoV-2 infection and the overlap between the two subgraphs. This overlap analysis allows for the identification of repurposing drugs targeting mechanisms that are modulated by viral infection.

**Figure 4:**
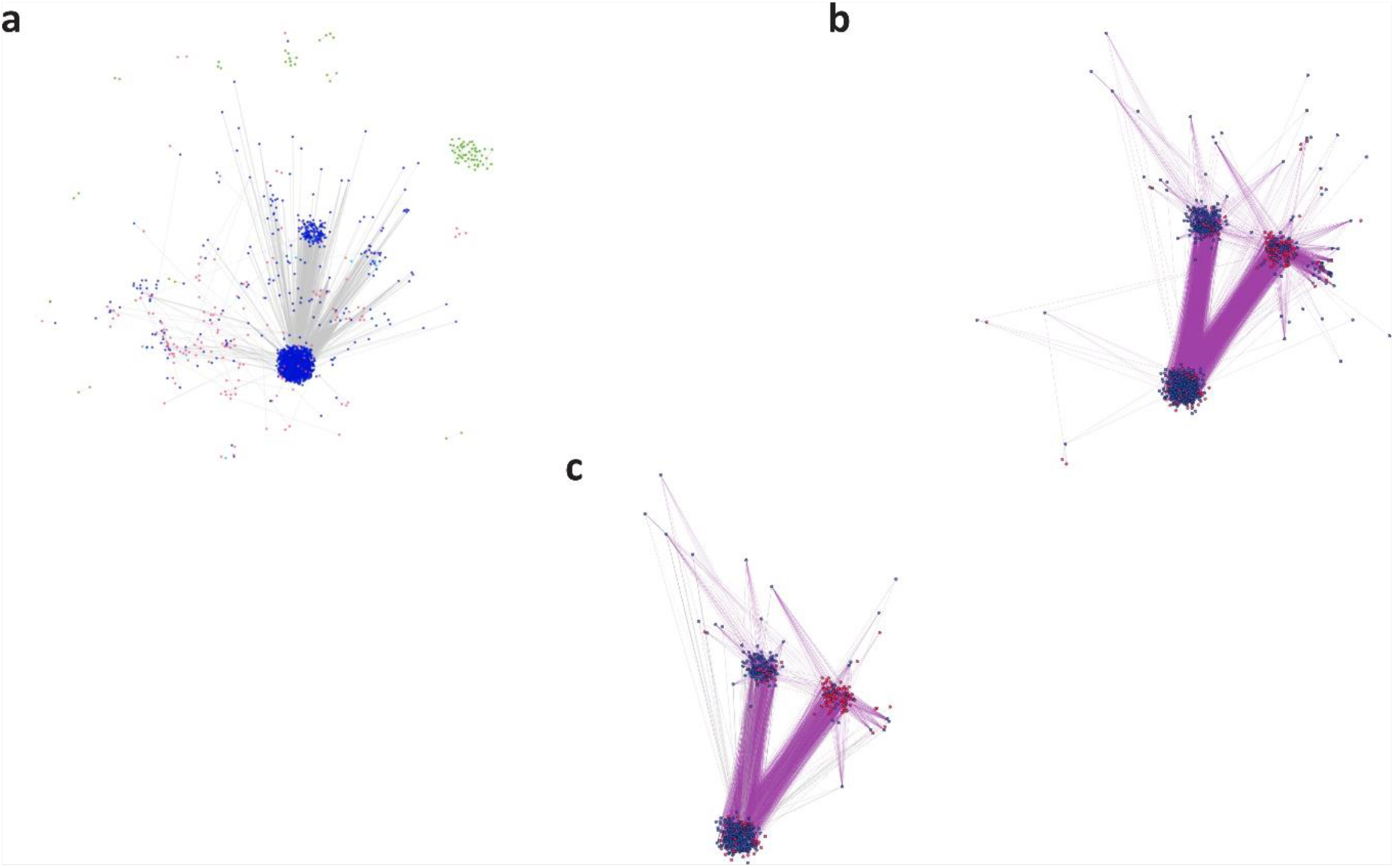
Identification of suitable targets for combination therapy by comparing subgraphs within the COVID-19 PHARMACOME. Incorporation of gene expression data into the COVID-19 PHARMACOME resulted in a subgraph characterized by the entities (genes/proteins) that respond to viral infection (a). Mapping of the filtered results obtained from drug repurposing screenings (IC50 < 10 μM) to the PHARMACOME resulted in a subgraph enriched for drug repurposing targets (b). The intersection between subgraphs presented in (a) and (b) is highly enriched for drug repurposing targets directly linked to the viral infection response (c).

A total number of 870 mechanisms were identified as being targeted by most of the drug repurposing candidates (see section “Associated pathway identification” in supplementary materials). When compared to the annotated subgraphs in the COVID-19 PHARMACOME, 201 of the 227 determined associated pathways found for the viral life cycle subgraph overlapped with those for the drug repurposing targets while the host response subgraph shared 90 of its 105 pathways.

### Mapping of drug repurposing signals to hypervariable regions of the COVID-19 PHARMACOME

One of the key questions arising from the network analysis is whether the repurposing drugs target mechanisms are specifically activated during viral infection. In order to establish this link, we mapped differential gene expression analyses from two single-cell sequencing studies to our COVID-19 PHARMACOME (see section “Differential Gene Expression” in supplementary material). An overlay of differential gene expression data (adjusted *p*-value ≤ 0.1 and abs(log fold-change) > 0.25) on the COVID-19 PHARMACOME reveals a distinct pattern characterized by the high responsiveness (expressed by variation of regulation of gene expression) to the viral infection (Figure 4A).

### Virus-response mechanisms are targets for repurposing drugs

In the next step, we analyzed which areas of the COVID-19 graph respond to SARS-CoV-2 infection (indicated by significant variance in gene expression) and are targets for repurposing drugs. To this end, we mapped signals from the drug repurposing screenings to the subgraph that showed responsiveness to SARS-CoV-2 infection (Figure 4B). Figure 4C depicts the resulting subgraph that is characterized by the transcriptional response to SARS-CoV-2 infection and the presence of target proteins of compounds that have been identified in drug repurposing screening experiments.

### The COVID-19 PHARMACOME supports rational targeting strategies for COVID-19 combination therapy

We mapped existing combinatorial therapy data to the COVID-19 PHARMACOME in order to evaluate its potential in guiding rational approaches towards combination therapy using repurposing drug candidates. Combinatorial treatment data obtained from the results published by Bobrowski et al.^40^ and Ellinger et al.^41^ were mapped to the COVID-19 PHARMACOME. Figure 5 provides an overview of the mapped compounds, thier protein targets, and the interaction mechanisms. Analysis of the overlaps between the drug repurposing screening data showed that four of the ten compounds reported in the synergistic treatment approach by drug repurposing data were represented in our initial non-redundant set of candidate repurposing drugs.

**Figure 5:**
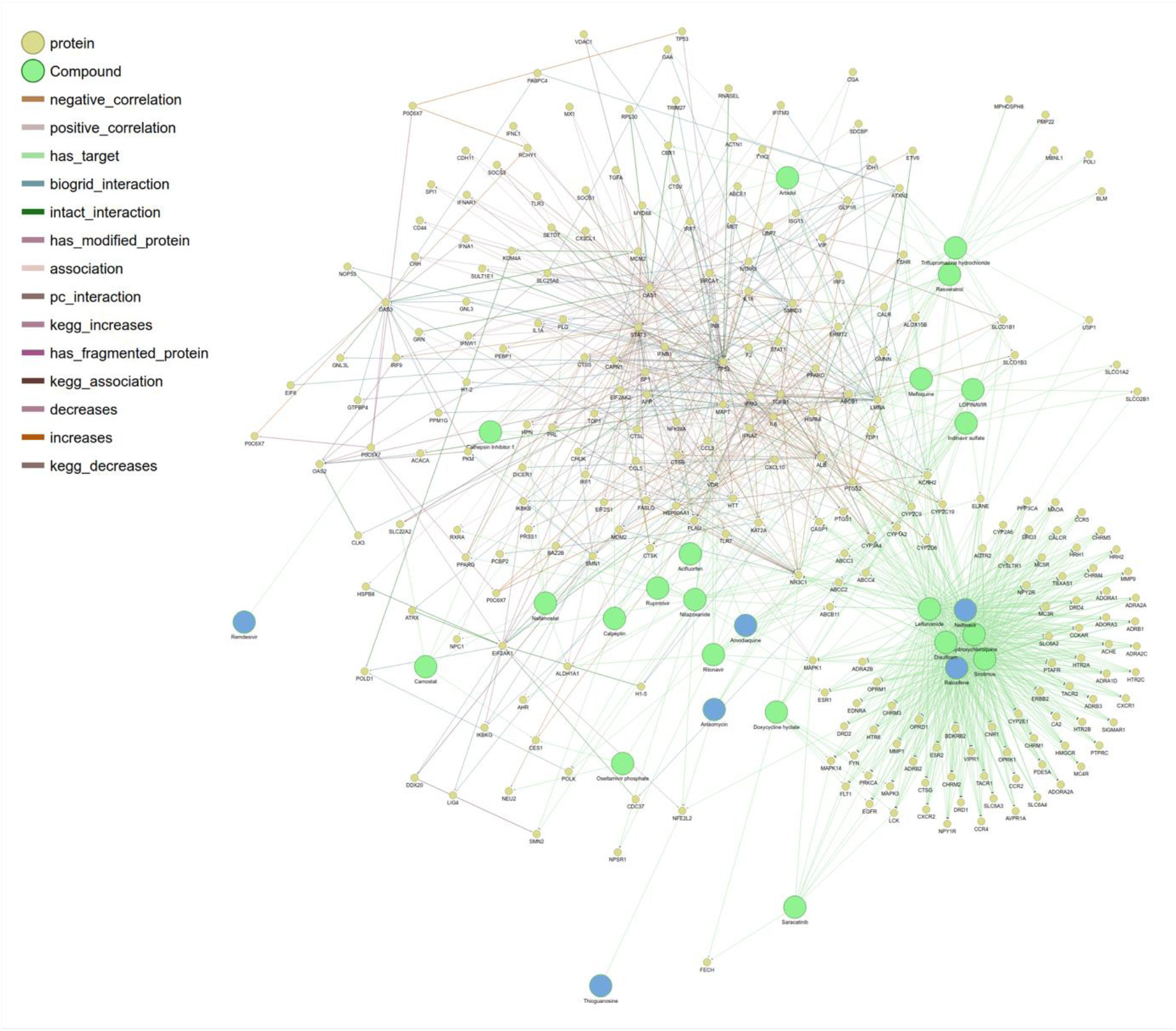
Visualization of drug repurposing candidates (and their targets) used in combination treatment experiments. The subgraph depicts the drug repurposing candidate molecules in relation to each other and their targets. Shortest path lengths between drug combinations were calculated from this subgraph and are available in the supplementary material (Supplementary Table 5).

Based on the association between repurposing drug candidates and the areas of the COVID-19 PHARMACOME that respond to SARS-CoV-2 infection (Figure 4), we hypothesized that the number of edges between a pair of drug nodes may be linked to the effectiveness of the drug combination (Supplementary Figure 2). In order to evaluate whether the determined outcome of a combination of drugs correlated with the distance between said drug nodes, we compared distances for combinations of drugs within the COVID-19 PHARMACOME for which their effect was known (Supplementary Tables 3 & 5). Of the 47 drug combinations we were able to check within the COVID-19 PHARMACOME, we found that the pairs of drugs known to have a synergistic effect in the treatment of SARS-CoV-2 had an average shortest path length of 2.43, while antagonistic combinations were found to be farther apart with an average shortest path length of 4.0 (Supplementary Table 7). Based on our calculations, we formulated three categories for predicting the outcome of new drug combinations on infection using the shortest path lengths between them within the COVID-19 PHARMACOME. Drug combinations with shortest path lengths of 2 indicate a synergistic relationship between the compounds, 3 was determined to be inconclusive as our calculations did not justify a specific outcome, and those with a shortest path length of 4 or more were predicted to have an antagonistic relationship.

In order to test our ability to predict the outcome of novel drug combinations, we selected five compounds: Remdesivir (a virus replicase inhibitor), Nelfinavir (a virus protease inhibitor), Raloxifene (a selective estrogen receptor modulator), Thioguanosine (a chemotherapy compound interfering with cell growth), and Anisomycin (a pleiotropic compound with several pharmacological activities, including inhibition of protein synthesis and nucleotide synthesis). These compounds were used in four different combinations (Remdesivir/Thioguanosine, Remdesivir/Raloxifene, Remdesivir/Anisomycin and Nelfinavir/Raloxifene) to test the potency of these drug pairings in phenotypic, cellular assays. Figure 6 shows the results of these combinatorial treatments on the virus-induced cytopathic effect in Caco-2 cells.

**Figure 6:**
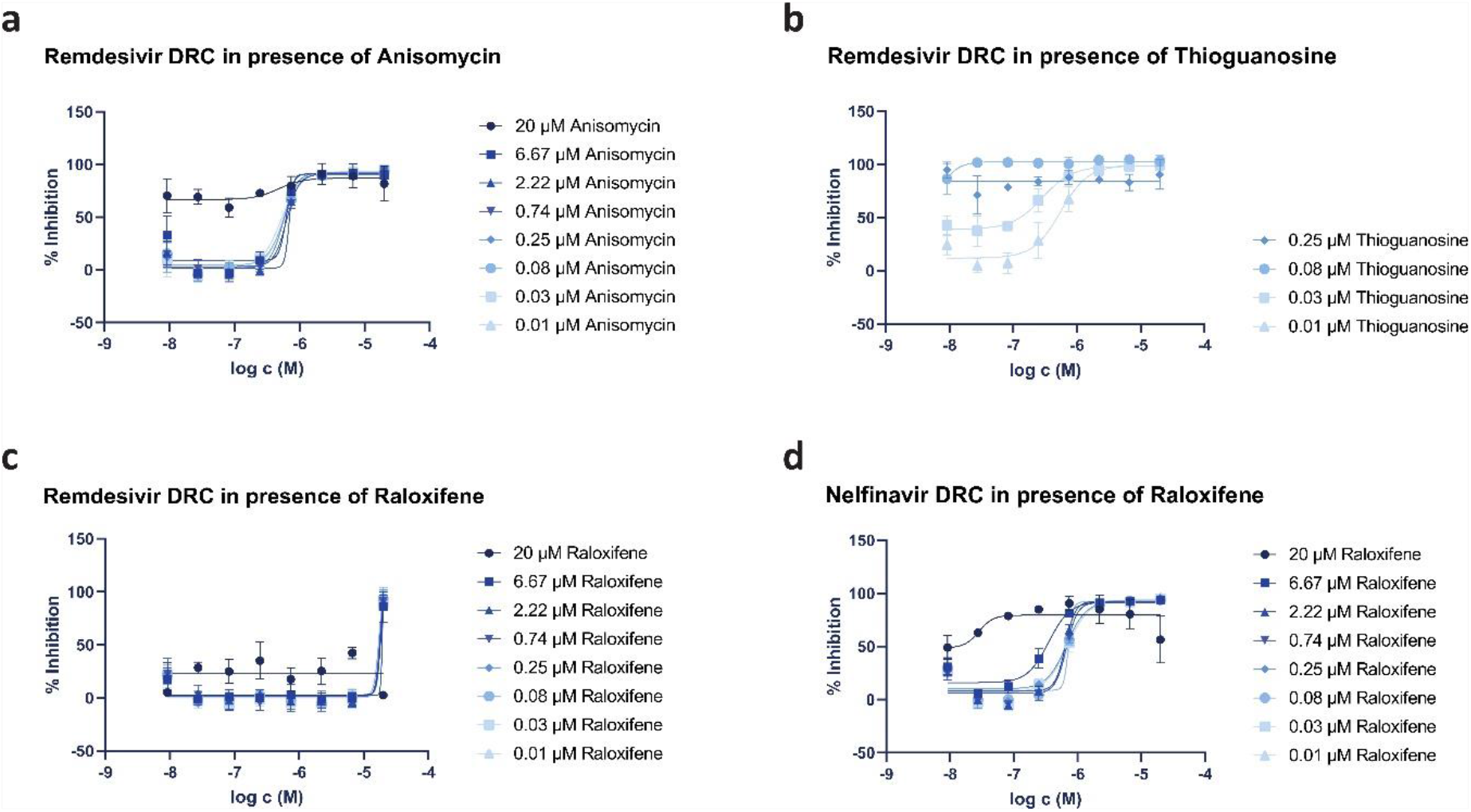
Dose-response curves (DRC) depicting viral inhibition of SARS-CoV-2 by select drug combinations. a) A threshold effect can be seen with the Remdesivir/Anisomycin combination when Anisomycin reaches 20 μM, well beyond Anisomycin’s IC50 alone. Remdesivir activity does not appear to be affected by Anisomycin, while Remdesivir seems to be equally affected (de-potentiated) by low to high concentrations of Raloxifene. b) Viral inhibition for Remdesivir/Thioguanosine can be seen only at lower Thioguanosine concentrations, at higher concentrations the clear curve shift of Remdesivir at lower concentration (effect beyond Loewe’s additivity formula) could not be appreciated. c) Raloxifene had an antagonistic effect on Remdesivir’s viral replication inhibition activity. d) A clear shift in Nelfinavir’s DRC can be observed when combined with Raloxifene, but also suggests a threshold effect when Raloxifene concentrations are higher than 2.2 μM.

Our results indicate that compound combinations acting on different viral mechanisms, such as Remdesivir and Thioguanosine (Figure 6b) or Nelfinavir and Raloxifene (Figure 6d), showed synergy, while compounds acting on host mechanisms, for instance Anisomycin or Raloxifene, when combined with Remdesivir (Figure 6a and Figure 6c, respectively), resulted in neither synergistic nor additive effects. Interestingly, our experiments revealed that the HIV-Protease inhibitor Nelfinavir, which already appeared to be active against viral post-entry fusion steps of both SARS-CoV^42^ and SARS-CoV-2^43^, displayed synergistic effects when combined with high concentrations of Raloxifene. This result agrees with our predictions generated using the COVID-19 PHARMACOME in which the drug combination with the shortest distance, Raloxifene and Nelfinavir (Supplementary Table 5), would have a synergistic effect on SARS-CoV-2 pathology.

## Discussion

By combining a significant number of knowledge graphs which represent various aspects of COVID-19 pathophysiology and drug-target information we were able to generate the COVID-19 PHARMACOME, a unique resource that covers a wide spectrum of cause-and-effect knowledge about SARS-CoV-2 and its interactions with the human host. Based on a systematic review of the results derived from published drug repurposing screening experiments, as well as our own drug repurposing screening results, we were able to identify mechanisms targeted by a variety of compounds showing virus inhibition in phenotypic, cellular assays. With the COVID-19 PHARMACOME, we are now able to link repurposing drugs, their targets and the mechanisms modulated by said drugs within one computable data structure, thereby enabling us to target - in a combinatorial treatment approach - different, independent mechanisms. By challenging the COVID-19 PHARMACOME with gene expression data, we have identified subgraphs that are responsive (at gene expression level) to virus infection. Network analysis along with the overview on previous repurposing experiments provided us with the insights needed to select the optimal repurposing drug candidates for combination therapy. Experimental verification showed that this systematic approach is valid; we were able to identify two drug-target-mechanism combinations that demonstrated synergistic action of the repurposed drugs targeting different mechanisms in combinatorial treatments.

We are fully aware of the fact that the COVID-19 PHARMACOME combines experimental results generated in different assay conditions. In the course of our work, we accumulated evidence that assay responses recorded using Vero E6 cells in comparison to Caco-2 cells may only partially overlap. Comparative analysis of the results of both assay systems to virus infection by means of transcriptome-wide gene expression analysis is one of the experiments we plan to perform next. However, for the identification of meaningful combinations of repurposing drugs, the current model-driven information fusion approach was shown to work well despite the putative differences between drug repurposing screening assays.

Given the urgent need for treatments that work in an acute infection situation, our approach described here paves the way for systematic and rational approaches towards combination therapy of SARS-CoV-2 infections. We want to encourage all our colleagues to make use of the COVID-19 PHARMACOME, improve it, and add useful information about pharmacological findings (e.g. from candidate repurposing drug combination screenings). In addition to vaccination and antibody therapy, (combination) treatment with small molecules remains one of the key therapeutic options for combatting COVID-19. The COVID-19 PHARMACOME will therefore be continuously improved and expanded to serve integrative approaches in anti-SARS-CoV-2 drug discovery and development.

## Acknowledgements

In part, this project is supported by the European Union’s Horizon 2020 research and innovation program under grant agreement No 101003551, project Exscalate4CoV.

## References

1 Xu, B., Gutierrez, B., Mekaru, S., Sewalk, K., Goodwin, L., Loskill, A., … & Zarebski, A. E. (2020). Epidemiological data from the COVID-19 outbreak, real-time case information. Scientific data, 7(1), 1–6.

2 Lipsitch, M., Swerdlow, D. L., & Finelli, L. (2020). Defining the epidemiology of Covid-19—studies needed. New England journal of medicine, 382(13), 1194–1196.

3 Holmdahl, I., & Buckee, C. (2020). Wrong but Useful—What Covid-19 Epidemiologic Models Can and Cannot Tell Us. New England Journal of Medicine.

4 Cao, W., & Li, T. (2020). COVID-19: towards understanding of pathogenesis. Cell Research, 1–3.

5 Liao, M., Liu, Y., Yuan, J., Wen, Y., Xu, G., Zhao, J., … & Liu, L. (2020). Single-cell landscape of bronchoalveolar immune cells in patients with COVID-19. Nature Medicine, 1–3.

6 Tay, M. Z., Poh, C. M., Rénia, L., MacAry, P. A., & Ng, L. F. (2020). The trinity of COVID-19: immunity, inflammation and intervention. Nature Reviews Immunology, 1–12.

7 Gervasoni, S.; Vistoli, G.; Talarico, C.; Manelfi, C.; Beccari, A.R.; Studer, G.; Tauriello, G.; Waterhouse, A.M.; Schwede, T.; Pedretti, A. A Comprehensive Mapping of the Druggable Cavities within the SARS-CoV-2 Therapeutically Relevant Proteins by Combining Pocket and Docking Searches as Implemented in Pockets 2.0. Int. J. Mol. Sci. 2020, 21, 5152.

8 Ostaszewski, M., Mazein, A., Gillespie, M. E., Kuperstein, I., Niarakis, A., Hermjakob, H., … & Schreiber, F. (2020). COVID-19 Disease Map, building a computational repository of SARS-CoV-2 virus-host interaction mechanisms. Scientific data, 7(1), 1–4.

9 Domingo-Fernandez, D. et al. COVID-19 Knowledge Graph: a computable, multi-modal, cause-and-effect knowledge model of COVID-19 pathophysiology. Bioinformatics. btaa834 (2020).

10 Gysi, D. M., Valle, Í. D., Zitnik, M., Ameli, A., Gan, X., Varol, O., … & Barabási, A. L. (2020). Network medicine framework for identifying drug repurposing opportunities for covid-19. arXiv preprint arXiv:2004.07229.

11 Khan, J. Y., Khondaker, M., Islam, T., Hoque, I. T., Al-Absi, H., Rahman, M. S., … & Rahman, M. S. (2020). COVID-19Base: A knowledgebase to explore biomedical entities related to COVID-19. arXiv preprint arXiv:2005.05954.

12 Kuperstein, I., Bonnet, E., Nguyen, H. A., Cohen, D., Viara, E., Grieco, L., … & Dutreix, M. (2015). Atlas of Cancer Signalling Network: a systems biology resource for integrative analysis of cancer data with Google Maps. Oncogenesis, 4(7), e160–e160.

13 Kodamullil, A. T., Younesi, E., Naz, M., Bagewadi, S., & Hofmann-Apitius, M. (2015). Computable cause-and-effect models of healthy and Alzheimer’s disease states and their mechanistic differential analysis. Alzheimer’s & Dementia, 11(11), 1329–1339.

14 Fujita, K. A., Ostaszewski, M., Matsuoka, Y., Ghosh, S., Glaab, E., Trefois, C., … & Diederich, N. (2014). Integrating pathways of Parkinson’s disease in a molecular interaction map. Molecular neurobiology, 49(1), 88–102.

15 Matsuoka, Y. et al. A comprehensive map of the influenza A virus replication cycle. BMC Syst. Biol. 7, 97 (2013

16 Khan, J. Y., Khondaker, M., Islam, T., Hoque, I. T., Al-Absi, H., Rahman, M. S., … & Rahman, M. S. (2020). COVID-19Base: A knowledgebase to explore biomedical entities related to COVID-19. arXiv preprint arXiv:2005.05954.

17 Ostaszewski, M., Mazein, A., Gillespie, M. E., Kuperstein, I., Niarakis, A., Hermjakob, H., … & Schreiber, F. (2020). COVID-19 Disease Map, building a computational repository of SARS-CoV-2 virus-host interaction mechanisms. Scientific data, 7(1), 1–4.

18 Blanco-Melo, D., Nilsson-Payant, B. E., Liu, W. C., Uhl, S., Hoagland, D., Møller, R., … & Wang, T. T. (2020). Imbalanced host response to SARS-CoV-2 drives development of COVID-19. Cell.

19 Gordon, D. E., Jang, G. M., Bouhaddou, M., Xu, J., Obernier, K., White, K. M., … & Tummino, T. A. (2020). A SARS-CoV-2 protein interaction map reveals targets for drug repurposing. Nature, 1–13.

20 Bojkova, D., Klann, K., Koch, B., Widera, M., Krause, D., Ciesek, S., … & Münch, C. (2020). Proteomics of SARS-CoV-2-infected host cells reveals therapy targets. Nature, 1–8.

21 Ashburn, T. T., & Thor, K. B. (2004). Drug repositioning: identifying and developing new uses for existing drugs. Nature reviews Drug discovery, 3(8), 673–683.

22 Pushpakom, S., Iorio, F., Eyers, P. A., Escott, K. J., Hopper, S., Wells, A., … & Norris, A. (2019). Drug repurposing: progress, challenges and recommendations. Nature reviews Drug discovery, 18(1), 41–58.

23 http://rdcu.be/qKdSKdSp://rdcu.be/qKdS

24 https://doi.org/10.1073/pnas.1810137115

25 https://reframedb.org/assays/A00461

26 https://reframedb.org/assays/A00440

27 preprint, DOI:.21203/rs.3.rs-23951/v1

28 Slater, T. (2014). Recent advances in modeling languages for pathway maps and computable biological networks. Drug discovery today, 19(2), 193–198.

29 Domingo-Fernández, D., Mubeen, S., Marín-Llaó, J., Hoyt, C. T., & Hofmann-Apitius, M. (2019). PathMe: Merging and exploring mechanistic pathway knowledge. BMC bioinformatics, 20(1), 243.

30 Domingo-Fernández, D., Hoyt, C. T., Bobis-Álvarez, C., Marín-Llaó, J., & Hofmann-Apitius, M. (2018). ComPath: an ecosystem for exploring, analyzing, and curating mappings across pathway databases. NPJ systems biology and applications, 4(1), 1–8.

31 Astghik, S. et al., submitted, Bioinformatics Journal (OUP)

32 Chua, R. L., Lukassen, S., Trump, S., Hennig, B. P., Wendisch, D., Pott, F., Debnath, O., Thürmann, L., Kurth, F., Völker, M.T., Kazmierski, J., Timmermann, B., Twardziok, S., Schneider, S., Machleidt, F., Müller-Redetzky, H., Maier, M., Krannich, A., Schmidt, S., Balzer, F., Liebig, J., Loske, J., Suttorp, N., Eils, J., Ishaque, N., Liebert, U.G., von Kalle, C., Witzenrath, M., Goffinet, C., Drosten, C., Laudi, S., Lehmann, I., Conrad, C., Sander, L-E. and Eils, R. (2020). COVID-19 severity correlates with airway epithelium–immune cell interactions identified by single-cell analysis. Nature Biotechnology, 38(8), 970–979.

33 Ravindra, N. G., Alfajaro, M. M., Gasque, V., Habet, V., Wei, J., Filler, R. B., Huston, N. C., Wan, H., Szigeti-Buck, K., Wang, B., Wang, G., Montgomery, R.R., Eisenbarth, S. C., Williams, A., Pyle, A.M., Iwasaki, A., Horvath, T.L., Foxman, E.F., Pierce, R.W., van Dijk, D., and Wilen, C.B. (2020). Single-cell longitudinal analysis of SARS-CoV-2 infection in human bronchial epithelial cells. bioRxiv.

34 Kuleshov MV, Jones MR, Rouillard AD, et al. Enrichr: a comprehensive gene set enrichment analysis web server 2016 update. Nucleic Acids Res. 2016;44(W1):W90–W97. doi:10.1093/nar/gkw377

35 https://pypi.org/project/gseapy/

36 Benjamini Y. Discovering the false discovery rate: False Discovery Rate. J. R. Stat. Soc. Ser. B Stat. Methodol. 2010;72(4):405–416. doi: 10.1111/j.1467-9868.2010.00746.x.

37 Hoehl, S., Rabenau, H., Berger, A., Kortenbusch, M., Cinatl, J., Bojkova, D., Behrens, P., Böddinghaus, B., Götsch, U., Naujoks, F., Neumann, P., Schork, J., Tiarks-Jungk, P., Walczok, A., Eickmann, M., Vehreschild, M., Kann, G., Wolf, T., Gottschalk, R., & Ciesek, S. (2020). Evidence of SARS-CoV-2 infection in returning travelers from Wuhan, China. New England Journal of Medicine, 382(13), 1278–1280.

38 Ellinger, B., Bojkova, D., Zaliani, A., Cinatl, J., Claussen, C., Westhaus, S., … & Gribbon, P. (2020). Identification of inhibitors of SARS-CoV-2 in-vitro cellular toxicity in human (Caco-2) cells using a large scale drug repurposing collection. manuscript under review

39 Bobrowski, T., Chen, L., Eastman, R. T., Itkin, Z., Shinn, P., Chen, C., Guo, H., Zheng, W., Michael, S., Simeonov, A., Hall, M., Zakharov, A.V., and Muratov, E.N. (2020). Discovery of Synergistic and Antagonistic Drug Combinations against SARS-CoV-2 In Vitro. BioRxiv.

40 Bobrowski, T., Chen, L., Eastman, R. T., Itkin, Z., Shinn, P., Chen, C., Guo, H., Zheng, W., Michael, S., Simeonov, A., Hall, M., Zakharov, A.V., and Muratov, E.N. (2020). Discovery of Synergistic and Antagonistic Drug Combinations against SARS-CoV-2 In Vitro. BioRxiv.

41 Ellinger, B et al. (2020). Identification of inhibitors of SARS-CoV-2 in-vitro cellular toxicity in human (Caco-2) cells using a large scale drug repurposing collection. Preprint. https://doi.org/10.21203/rs.3.rs-23951/v1.

42 Yamamoto, N., Yang, R., Yoshinaka, Y., Amari, S., Nakano, T., Cinatl, J., … & Tamamura, H. (2004). HIV protease inhibitor nelfinavir inhibits replication of SARS-associated coronavirus. Biochemical and biophysical research communications, 318(3), 719–725.

43 Musarrat, F., Chouljenko, V., Dahal, A., Nabi, R., Chouljenko, T., Jois, S. D., & Kousoulas, K. G. (2020). The anti-HIV Drug Nelfinavir Mesylate (Viracept) is a Potent Inhibitor of Cell Fusion Caused by the SARS-CoV-2 Spike (S) Glycoprotein Warranting further Evaluation as an Antiviral against COVID-19 infections. Journal of medical virology.

