## Supplementary Material for "The COVID-19 PHARMACOME: A method for the rational selection of drug repurposing candidates from multimodal knowledge harmonization"

| Disease Map Name | Reference | Abbreviation | Source of Information | Modalities & Scales Represented | Model scope |
|---|---|---|---|---|---|
| Interactome | Gordon, D. E., Jang, G. M., Bouhaddou, M., Xu, J., Obernier, K., White, K. M., ... & Tummino, T. A. (2020). A SARS-CoV-2 protein interaction map reveals targets for drug repurposing. *Nature*, 1-13. | gordon | Pull-down experiments | Molecular entities | Host virus interaction |
| COVID-19 Disease Map | Ostaszewski, M., Mazein, A., Gillespie, M. E., Kuperstein, I., Niarakis, A., Hermjakob, H., ... & Schreiber, F. (2020). COVID-19 Disease Map, building a computational repository of SARS-CoV-2 virus-host interaction mechanisms. *Scientific data*, *7*(1), 1-4. | lux | Literature mining and manual curation | Pathway level<br><br>Pathway model | Host virus interaction and pathways |
| COVID-19 Knowledge Graph | Domingo-Fernandez, D., Baksi, S., Schultz, B., Gadiya, Y., Karki, R., Raschka, T., ... & Hofmann-Apitius, M. (2020). COVID-19 Knowledge Graph: a computable, multi-modal, cause-and-effect knowledge model of COVID-19 pathophysiology. *BioRxiv*. | cbm_scai | Literature mining and manual curation | Multimodal and multiscale, spanning from molecular entities to clinical phenotypes<br><br>Cause-and-effect model | Host virus interactions and mechanisms |
| COVID-19 Interaction Graph (Elsevier) | https://pharma.elsevier.com/covid-19/elsevier-models-for-covid19-bio-molecular-mechanisms/ | elsevier | Literature mining | Molecular entities | Host virus interaction |
| COVID-19 Sepsis Risk Factor Graph Model | https://precisionlife.com/wp-content/uploads/2020/05/precisionlife-Sepsis-COVID- | | Driven by genetics data analysis; combining data-driven and | Molecular scale, disease phenotype focus on sepsis | Focus on sepsis biology and virus susceptibility |

| | | | | | |
|---|---|---|---|---|---|
| | 19-Risk-Factors-Report.pdf | | knowledge-driven approaches | | |
| COVID-19 Proteome | Bojkova, D., Klann, K., Koch, B., Widera, M., Krause, D., Ciesek, S., ... & Münch, C. (2020). Proteomics of SARS-CoV-2-infected host cells reveals therapy targets. *Nature*, 1-8. | | Driven by proteomics analysis; | Molecular scale, comparative approach with infected / non-infected cells | Virus host interaction |
| COVID-19 and Lung Epithelial Cell Models | Schlage, W. K., Westra, J. W., Gebel, S., Catlett, N. L., Mathis, C., Frushour, B. P., ... & Lietz, M. (2011). A computable cellular stress network model for non-diseased pulmonary and cardiovascular tissue. *BMC systems biology*, *5*(1), 168.<br><br>Park, J. S., Schlage, W. K., Frushour, B. P., Talikka, M., Toedter, G., Gebel, S., ... & Kogel, U. (2013). Construction of a computable network model of tissue repair and angiogenesis in the lung. *J Clinic Toxicol S*, *12*, 2161-0495. | pmi | Driven by literature analysis and expert knowledge | Molecular level; tissue specific (lung epithelium and microvasculature) | Target cell physiology |
| Heme Knowledge Graph | Humayun, F., et al. (2020). A computational approach for mapping heme biology in the context of hemolytic disorders. Frontiers in Bioengineering and Biotechnology, 8, 74. | heme | Manual curation | Molecular entities | Heme physiology including major blood - based physiology |
| BioDati COVID-19 Model | https://networkstore.demo.biodati.com/networks/01E46GDFQAGK5W8EFS9S9WMH12?format=normal | biodati | Literature mining | Molecular entities | Related molecular interactions |
| IntAct Coronavirus Molecular Interaction Dataset | https://www.ebi.ac.uk/intact/query/annot:%22dataset:coronavirus%22 | intact | Literature Mining | Molecular entities | Related molecular interactions |

**Supplementary Table 1**. Constituent COVID-19 models that were incorporated into the COVID-19 supergraph.

| Authors | Title | URL | Library Screened and Number of Hits | Cell Type Tested |
|---|---|---|---|---|
| Touret et al. | In vitro screening of a FDA approved chemical library reveals potential inhibitors 1of SARS-CoV-2 replication | https://www.biorxiv.org/content/10.1101/2020.04.03.023846v1 | 6 hits<br><br>Prestwick Chemical Library® (a library of 1,520 off-patent small molecules) | VeroE6 (ATCC CRL-1586) cells |
| Gordon et al. | A SARS-CoV-2 protein interaction map reveals targets for drug repurposing | https://www.nature.com/articles/s41586-020-2286-9 | 36 hits out of 75 pre-selected compounds | VeroE6 (ATCC CRL-1586) cells |
| Riva et al. | A Large-scale Drug Repositioning Survey for SARS-CoV-2 Antivirals | https://www.nature.com/articles/s41586-020-2577-1 | ReFRAME library (approx. 12,000 compounds), 18 hits | VeroE6 (ATCC CRL-1586) cells |
| Jeon et al. | Identification of antiviral drug candidates against SARS-CoV-2 from FDA-approved drugs | https://aac.asm.org/content/64/7/e00819-20.abstract | Initial screening of approx. 3000 cmpd with MERS / SARS-CoV-1; re-screening of 48 drugs on SARS-CoV-2; 24 hits identified | Vero (ATCC CCL-81) cells |
| Weston et al. | Broad anti-coronaviral activity of FDA approved drugs against SARS-CoV-2 in vitro and SARS-CoV in vivo | https://www.biorxiv.org/content/10.1101/2020.03.25.008482v2.full.pdf | Targeted screening of pre-selected 20 compounds; 17 hits | VeroE6 (ATCC CRL-1586) cells |
| Ellinger et al. | Identification of inhibitors of SARS-CoV-2 in-vitro cellular toxicity in human (Caco-2) cells using a large scale drug repurposing collection | https://www.researchsquare.com/article/rs-23951/v1 | 5632 compounds (Fraunhofer replica Broad library); 77 hits | Human Caco–2 cells |

**Supplementary Table 2.** Overview on published and proprietary drug repurposing data used in this study.

**Drug repurposing screening using phenotypic assays**

A highly qualified set of known bioactives and marketed compounds (the Fraunhofer Repurposing Collection, established as a mirror set using principles set up by the Broad Institute [i] [ii]) with a well-defined collection of 5632 compounds including 3488 that have undergone previous clinical investigations (approved drugs, phases I–III, and withdrawn compounds) across 600 indications is one of the most comprehensive sets of annotated compounds currently described, with 5682 unique compounds. The collection includes 3,400 compounds that have reached clinical use across 600 indications. In addition, the collection contains 1582 pre-clinical compounds at varying stages of validation. In 2019, the compounds were purchased from the same set of more than 70 high-quality suppliers identified by the Broad Institute and were quality controlled by LC/MS for purity and identity (minimum purity > 90%). The compounds were stored at a concentration of 10 mM in 100% DMSO at -20 °C. A curated database is available listing the compounds, indications, primary targets (where known), and mechanism of action, as well as analysis tools which can help to determine the mechanism of action and target. This collection of compounds differs from many in that it contains a high proportion of clinical and preclinical candidates, as well as marketed drugs which are commonly found in classical repurposing collections. The collection has been screened in phenotypic antiviral assays, either in epithelial cells, human (Caco-2) or cells derived from green macaque (Vero-E6). The cells were treated with or without virus and the cytotoxic effect of each molecule was measured after 48 hr (Caco-2) or 120 hr (Vero-E6) post infection.

**Gene expression data analysis**

To validate the edges in the supergraph, differential expression data were obtained from two studies that generated single-cell RNA sequencing (scRNA-seq) data for SARS-CoV-2 infected samples.

The first data set is comprised of the results taken from Ravindra et al.[iii]. Primary human bronchial epithelial cells (HBECs) were cultured for 28 days prior to SARS-CoV-2 infection. Cells were kept in culture for three days; each day a suspension was taken and prepared for single-cell RNA sequencing. Using the 10X Genomics cellranger pipeline, expression reads were mapped against the human and viral genomes and count matrices were generated. Then, the Seurat[iv] package was used for clustering, and the cells were annotated based on marker genes reported in the molecular cell atlas[v]. Subsequently, cells were classified as infected if more than ten viral transcript counts were found. Finally, differential gene expression analysis was conducted; the authors pooled the three time-points and compared infected versus bystander, infected versus mock, and bystander versus mock. For a detailed description of their methodology, see Ravindra et al. The results of the differential gene expression analyses were retrieved from the Van Dijk GitHub repository[vi].

The second is the data set from Chua et al.[vii]. The authors conducted an observational cohort study at the Berlin and Leipzig university hospitals and acquired scRNA-seq data for 19 patients with moderate or critical disease and five healthy controls. Nasopharyngeal and bronchial specimens were taken and prepared for 3' single-cell RNA-sequencing. The raw data were processed with the 10X Genomics Cell Ranger[viii] [ix] and the Seurat package. The variables “sex” and “days past the first symptoms” have been used as confounder variables (cfr. Chua et al. for further details on the sample acquisition and data processing).

In both cases, the list of differentially expressed genes was used to validate that the experimental data agrees with the information from the supergraph. For every edge of the graph, it was tested if the experimental data either has supporting, contradicting, or no information about the involved genes.

**Graph visualization and layout**

Images of the COVID-19 PHARMACOME and its subgraphs were generated using Gephi[x], an open-source software created for graph and network analysis. The graph layout method used for visualization is a derivation of the Fruchterman-Reingold[xi] algorithm called OpenOrd[xii]. This algorithm that was modified to better handle larger graphs (greater than 1000 nodes) while still maintaining the ability to accurately distinguish clusters through a combination of simulated annealing iterations and edge cutting.

Relative node abundance and interrelationship images were created using the graph visualization software Cytoscape[xiii]. Nodes were labeled and colored by type such as protein, pathology, or other biological concepts and entities. Nodes whose types comprised less than 1% of total nodes in the COVID-19 PHARMACOME were not included in the final visualizations in order to improve image quality. Clusters were subsequently generated for each node type with the size of cluster being directly proportional to the number of nodes belonging to that class. Edges between individual nodes are also shown, but the total number of edges between any two clusters of node classes was limited to 1000 in order to ensure proper visualization. Clusters were arranged manually to optimize visibility of edges.

### Model overlap analysis

The COVID-19 supergraph consists of 10 constituent graph models that have been unified and harmonized using the OpenBEL language. For comparison, node entity values were extracted from individual models and, in the case of nodes representing genes, RNA, proteins, or any combination thereof, mapped to their UniProt accession numbers. The intersections of these sets were compared using Venn diagrams and the resulting overlapping values were analyzed.

### Rational selection of repurposing drugs for combination treatment

Currently, there are two experiments known to us that use combinations of (repurposing) drugs in phenotypic assays: the publication by Bobrowski et al.[xiv] and the work published by Ellinger et al.[xv]. Initially, we determined to what extent the compounds used by Bobrowski et al. are represented in the drug repurposing experiments performed by Ellinger et al. To this end, we identified all compounds and their targets from both manuscripts using the ChEMBL database, however, in some cases we had to manually select drug targets from other sources such as [xvi] or [xvii]. These compounds and targets were subsequently mapped to the COVID-19 supergraph.

A complete overview of the compound pairs and their outcomes used in both experiments is provided in Supplementary Tables 3 & 4. Supplementary Table 3 contains all published combinations from Bobrowski et al. as well as the four combinations described here. Drug combinations for which target/pathways could be retrieved from ChEMBL, the relative pathway overlap was calculated. For these pairs of drugs, compounds with a higher number of targets/pathways are combined into the primary combination compound (Compound 1). Detailed information on individual drug activities and path length calculations can be found in Supplementary Tables 4-7. It is important to note that for some combinations of drugs tested, such as Remdesivir and Anisomycin, there exists a threshold effect for which the presence of a second compound does not continuously modulate the effect of the first.

**Supplementary Table 3**: Overview of all drug combinations and their outcomes. The "overlap_pathways_relative" columns were calculated by dividing that row's "overlap_pathways" value by the corresponding "num_pathways_gsea" value. The color gradients go from white to a darker shade where the darker the shade, the closer the value is to the maximum of the column.

**Supplementary Table 4**: Individual drug activities by cell type.

**Supplementary Table 5**: The average shortest path length, number of associated pathways, and determined effect on SARS-CoV-2 infection for drug combinations tested.

**Supplementary Table 6**: Cytopathic effect (CPE) concentrations for each individual drug repurposing candidate.

**Supplementary Table 7**: Overview of the shortest path lengths and outcome averages.

**Supplementary Figure 1**: Graphical depiction representing the distribution of major classes of entities (pathologies, molecular entities such as proteins, drugs) in the COVID-19 PHARMACOME.

**Supplementary Figure 2**: Workflow representation for identifying new combination therapy drug candidates through multimodal modelling and path length calculation. a) Specific mechanisms (blue lines) affected by infection, referred to as hypervariable (HV) regions, are identified in the COVID-19 PHARMACOME and the involved components are marked as blue circles. b) Repurposing drug candidate hits (red circles) extracted from screenings are mapped to targets found in (a) (red lines). c) The shortest paths (purple lines) between known drug combinations are analyzed and the lengths of these paths are calculated. Here, shorter lengths are found to correlate with drug combinations that synergize while longer path lengths with those that are antagonistic. d) Determined associations between shortest path length and combination drug therapy outcome are used to predict new drug combinations in the COVID-19 PHARMACOME (orange lines).

---

[i] https://www.nature.com/articles/nm.4306
[ii] https://www.broadinstitute.org/drug-repurposing-hub
[iii] https://doi.org/10.1101/2020.05.06.081695
[iv] https://doi.org/10.1016/j.cell.2019.05.031
[v] https://doi.org/10.1101/742320
[vi] https://github.com/vandijklab/HBEC_SARS-CoV-2_scRNA-seq
[vii] https://www.nature.com/articles/s41587-020-0602-4
[viii] https://doi.org/10.1101/303727
[ix] https://doi.org/10.1186/s13059-015-0844-5
[x] Bastian M., Heymann S., Jacomy M. (2009). Gephi: an open source software for exploring and manipulating networks. International AAAI Conference on Weblogs and Social Media.
[xi] Fruchterman, T. M. J., & Reingold, E. M. (1991). Graph Drawing by Force-Directed Placement. Software: Practice and Experience, 21(11).
[xii] S. Martin, W. M. Brown, R. Klavans, and K. Boyack, “OpenOrd: An Open-Source Toolbox for Large Graph Layout,” SPIE Conference on Visualization and Data Analysis (VDA)., 2011
[xiii] Michael E. Smoot, Keiichiro Ono, Johannes Ruscheinski, Peng-Liang Wang, Trey Ideker, Cytoscape 2.8: new features for data integration and network visualization, Bioinformatics, Volume 27, Issue 3, 1 February 2011, Pages 431–432
[xiv] Bobrowski, T., Chen, L., Eastman, R. T., Itkin, Z., Shinn, P., Chen, C., Guo, H., Zheng, W., Michael, S., Simeonov, A., Hall, M., Zakharov, A.V., and Muratov, E.N. (2020). Discovery of Synergistic and Antagonistic Drug Combinations against SARS-CoV-2 In Vitro. BioRxiv.
[xv] Ellinger, B et al. (2020). Identification of inhibitors of SARS-CoV-2 in-vitro cellular toxicity in human (Caco-2) cells using a large scale drug repurposing collection. Preprint. https://doi.org/10.21203/rs.3.rs-23951/v1.
[xvi] https://www.drugbank.ca/
[xvii] https://www.probes-drugs.org/home/
