## Supplementary figures and images for "The COVID-19 PHARMACOME: A method for the rational selection of drug repurposing candidates from multimodal knowledge harmonization"

### Supplementary Figure 1

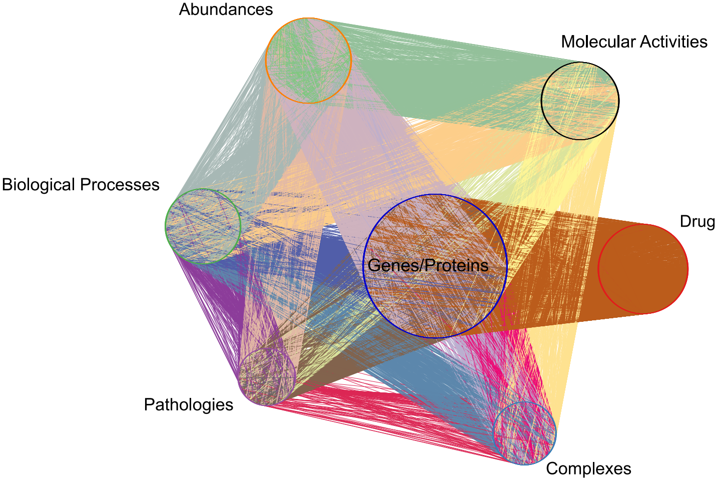

### Supplementary Figure 2

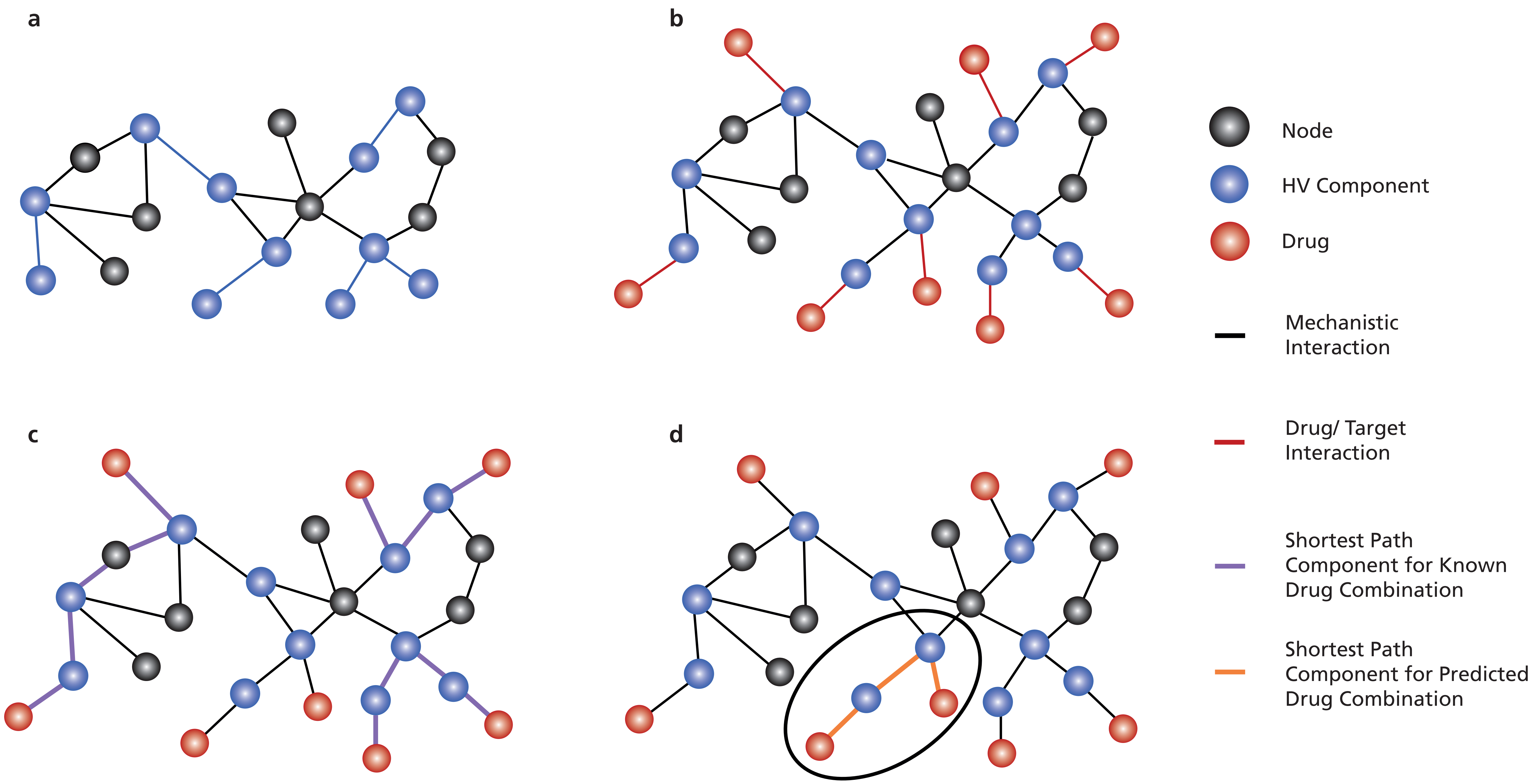
